# EXPLORING SOCIAL COGNITION DECLINE ACROSS ALZHEIMER’S DISEASE SEVERITY: INSIGHTS FROM A COMPARATIVE ANALYSIS

**DOI:** 10.1101/2024.12.17.629002

**Authors:** Gabriela Carneiro Martins, Vera Lucia Duarte Vieira, Flavia Miluzzi Pinetti, Ana Maria Leme de Arruda, Daniela Maruta Siqueira, Isadora Salvador Rocco, Paulo Henrique de Oliveira Bertolucci, Sergio Tufik, Claudia Berlim de Mello

## Abstract

The detection of subtle behavioral signs in early stages of Alzheimer’s Disease (AD) is of great importance to start clinical investigation. Memory and reasoning problems are usually more commonly reported by relatives than those associated with Social Cognition (SC). This study focusing on deficits in SC as a behavioral marker of Alzheimer’s Disease (AD) investigated: (1) selective impairments in SC domains according to stage of disease; (2) if impairments were better explained by nonsocial cognitive dysfunctions. Sixty-five elderly distributed into groups according to the Mini–Mental State Examination (Healthy, Mild cognitive impairment, Mild-AD, Moderate-AD) participated in the study. All underwent the Edinburgh Social Cognition Test (ESCoT), which investigates the cognitive and affective domains of Theory of Mind, as well as interpersonal and intrapersonal understanding of social norms. The nonsocial cognitive battery included traditional tests of episodic memory, executive functions, verbal fluency and language comprehension. Mediation analysis was performed to understand the influence of nonsocial cognitive skills in SC impairment among groups. Results revealed prominent losses in SC in moderate AD regardless of memory- skills. Theory of mind was a crucial link between social and nonsocial cognitive abilities. Executive functioning, verbal fluency and language comprehension mediated group-related cognitive decline’s effects on SC impairment. These findings underscore the complex interplay between cognitive domains in AD and suggest that selective impairments of SC highlight is as potential a marker for tracking the progression of the disease.

## INTRODUCTION

Alzheimer’s Disease (AD) is the most prevalent dementia syndrome in pathological aging, accounting for 60-70% of cases¹. The condition is characterized by a marked cognitive decline that over time results in loss of autonomy and quality of life. In a pre-clinical stage, cognitive complaints identified as more intense than expected for the individual’s age and educational level but that do not interfere with daily activities may characterize Mild Cognitive Impairment – MCI².

Efforts have been made worldwide for early detection of the disease. Biochemical diagnosis based on detection of biological markers such as Tau protein, phosphorylated Tau and beta-amyloid peptide in cerebrospinal fluid (CSF) is problematic as this is an invasive exam. Cognitive markers are the main evidence in initial stages. Impairments in episodic memory and executive functions, including cognitive inhibition and attentional control, are most evident³. Deficits in social cognition have also been reported and may be expressed as increasing loneliness or a trend for negative interactions^4^.

Social Cognition (SC) refers to a set of mental processes that regulate an individual’s behavior in the context of social interactions^5^. It encompasses different domains including Theory of Mind (ToM), the ability to understand that other people have intentions, desires and beliefs of their own and therefore may predict their behaviors^6^. Two subdomains of ToM have been proposed: Cognitive ToM (cToM), the ability to infer thoughts and beliefs, and Affective ToM (aToM), the ability to infer emotions and feelings^7^. Understanding social norms as a component of SC has been investigated^8^. Social norms have been defined as customary rules that define behaviors as appropriate or not^9^. Thus, the comprehension of social norms, in an interpersonal or intrapersonal perspective, is essential for social relationships.

Studies dedicated to investigate the extent to, and what components of, CS are affected along aging have revealed contradictory findings, in healthy elderly^2,8,10^ and in AD individuals^11,12^. The complexity of neuropsychological testing and the influence of global cognitive impairments, such as in attention or language, on task performance are some of the challenges.

A new multidimensional procedure developed by Baksh et al^8^, the Edinburgh Social Cognition Test (ESCOT), may be especially helpful to overcome methodological challenges. It assesses four components of CS: affective and cognitive ToM, interpersonal and intrapersonal understanding of social norms. Because all skills are investigated in a single set of stimuli, animated vignettes expressing characters interacting in familiar social scenes, it may be a less exhaustive and less cognitively demanding alternative. The authors evaluated young (mean 26.20 years), middle-aged (mean 50.60 years), and older (mean 72.45 years) healthy adults and found an effect of age on the cToM and aToM tasks. Progressive losses with increasing age were observed only in the ToM components of the test. These results suggest therefore that social norm comprehension skills remain preserved even in older ages.

This study aimed to investigate selective impairments in ToM and understanding of social norms according to severity of AD, and if performance would be influenced by memory and executive functions. To this end we compared individuals at different stages of AD (mild and moderate), to healthy controls (HC) and individuals with Mild Cognitive Impairments (MCI).

We hypothesized that (a) selective deficits in domains of ToM and understating of social norms would be observed in individuals with distinct clinical presentations, (b) such selective deficits would be found in mild AD, while a global impairment would predominate in the moderate stage; (c) SC deficits would not be explained by the non-social cognitive functions.

Identifying such specific and selective impairments in different clinical manifestations may elucidate the contribution of the role of SC as a behavioral mark of severity of dementia. Understanding specifically and selectively social cognition dysfunctions in different stages of dementia can strengthen early diagnosis and drive rehabilitation management of patients.

## MATERIAL AND METHODS

### Participants

Participants in the clinical groups were recruited from Outpatient Clinics of reference centers in São Paulo: S*erviço de Avaliação e Reabilitação do Idoso* (Associação Fundo de Incentivo à Pesquisa) and *Ambulatório de Neurologia Comportamental* do Hospital São Paulo (Universidade Federal de São Paulo). Diagnoses were established by medical specialists and confirmed by neuropsychological assessment according to DSM-5 ^13^ HC participants were outreached by social networks. All participants and, for AD participants also relatives, signed informed consents before data collection.

A sample of 65 elderly, 60 to 91 years-old (74.7± 8.17), 75.3% female, participated in this exploratory study with a cross-sectional design. They were distributed in four groups, according to level of cognitive impairments: (1) HC group, composed by individuals without cognitive complaints (n=17); (2) MCI group (n=17), (3) patients diagnosed with early-stage AD (n=15); (4) patients with moderate-stage AD (n=16). The classification of dementia groups was based on scores obtained in the Brazilian version of the Mini-Mental State Examination – MMSE ^14^.

Inclusion criteria for all groups were: to have reached at least four years of formal education, older than 60 years e have been submitted to clinical and neuropsychological assessment for diagnostic and definition of clinical presentation. Exclusion criteria were presence of (a) clinical indicators of depression (more than 10 points out of 30), according to the Brazilian versions of the Geriatric Depression Scale (GDS) ^15^, or anxiety (more than 10 points out of 30), according to the Geriatric Anxiety Inventory (GAI) ^16^; (b) previous neurological conditions associated with major illness (e.g. history of stroke, mixed dementia); (c) uncontrolled chronic diseases (e.g. diabetes, thyroid dysfunction); (c) uncorrected visual deficits.

### Measures

In the assessment of broad and non-social cognitive functions we emphasized language, episodic memory, executive functions and working memory. For this purpose, we adopted neuropsychological tests traditionally used in clinical practice, prioritizing those validated for the Brazilian population. Language skills were assessed using the Comprehension subtest of the Brazilian version of the Wechsler Adult Intelligence Scale (WAIS-III) ^18^. Episodic memory was assessed by means of Brazilian alternative versions of the logical story subtest of the Wechsler Memory Scale ^19^. Additionally we used the phonological (FAS) and semantic (animals) verbal fluency tests as a measure of lexical access ^20^. Finally, for the assessment of executive functions we used the Clock Drawing test (CDT)^21^. Working memory was assessed using digit span backwards. In all these tasks, performance was evaluated considering the total correct answers.

To assess Social Cognition skills, the Edinburgh Social Cognition Test (ESCOT) ^8^ was used, with the authors’ permission. The ESCOT investigates four social cognitive skills: cognitive ToM (cTOM); affective ToM (aTOM); interpersonal understanding of social norms and intrapersonal understanding of social norms. ESCOT consists of short animations involving cartoon-style drawings that express dynamic social interactions, in different contexts. There are five scenarios of interactions involving violation of social norms and five interactions that do not. Each animation lasts 30 seconds. At the end, a static image depicting a summarized version of the interaction is presented, which remains on the screen during the trial to avoid working memory demands. Participants were asked to describe what occurred in the interaction. They then answered questions from each of the four social cognition domains.

The ESCOT test consists of five videos of interactions considered as socially appropriate, since the character performs a socially appropriate action, and five videos with interactions considered as inappropriate. One question was to analyze possible effects of the type of social appropriateness (positive or negative) on the participants’ performance according to the degree of severity. Comparative analyses were made of the performance of each group of participants, with reference to that of the HC ones, in each domain of social cognition (ToM, understanding of social norms, normal) considering the subsets of videos considered to involve socially appropriate (n=5 videos) and socially inappropriate (n=5 videos) situations.

The cToM question asks "what is the character thinking?" The aToM question asks "how does the character feel at the end of the animation?" The question on interpersonal understanding of social norms asks "Did the other character in the scene of the animation behave as other people should behave?" Finally, the question on intrapersonal understanding of social norms asks whether the participant would act in the same way as the character. When the answer was limited, they were asked to respond to the following question: "Can you say more about what you mean by that? " or "can you explain this in a little more detail?" The total score for the whole test is 120.

### Procedures

The study was conducted during pandemic years (2021-2022) and therefore volunteers were assessed through video calls, in attention to the recommendations for physical distancing. The first meeting was dedicated to clinical interviews. HC participants (or relatives, usually daughters, for those from MCI and AD groups) were interviewed to obtain sociodemographic and clinical data. Additionally, they answered the Clinical Dementia Rating - CDR ^17^ to characterize current dementia symptoms.

Neuropsychological assessment was performed in two 60 minute sessions, one for diagnostic confirmation and characterization of global neuropsychological functioning, and the other for assessment of social cognition skills. For diagnostic confirmation it was used the Brazilian version of the Mini-Mental State Examination ^14^ with all participants. Healthy elderly adults who were familiar with computer use completed the neuropsychological tests independently. Participants from clinical groups were followed by a relative, in order to stimulate participant attentional focus in the tasks and to help in handling materials. The relative was previously explained about global procedures and oriented how to help participants and to avoid giving any clues for correct answers.

### Statistical Analysis

Sample size was calculated using the GPower software, using an effect size of 0.64 based on values found by Fliss et al ^12^ in a task with a purpose similar to the present study. A sample size of 32 participants was suggested for one-way ANOVA of 4 groups with a Beta=0.20 and alpha=0.05. Categorical data were presented as relative and absolute frequencies, and compared among groups with Fisher’s exact test. Shapiro-Wilk test was used to investigate data with Gaussian distribution. Normally distributed data were presented as mean and standard deviation and compared among groups with analysis of variance (ANOVA). Those non-normally distributed presented as median interquartile range and treated with non-parametric tests (Kruskal Wallis).

Generalized Linear Models (GzLM) were applied to investigate the effect of groups and confounders over the SC tests, allowing for the use of others than gaussian distribution. The optimal distribution of the dependent variable of models was chosen by the lower AIC, and residuals were tested with a Q-Q plot and Shapiro Wilk to ensure the quality of modeling with normally distributed error. The Bonferroni post hoc test was used to perform pairwise comparisons between groups.

Confirmatory Factor Analysis (CFA) was performed to resume main cognitive skills: episodic memory, working memory, verbal fluency, verbal comprehension and executive functions. A Principal Component Analysis was performed to enable the output of factors for the investigation in network and mediation analyses, respecting the criteria of Kaiser-Meyer-Olkin (KMO) test for sampling adequacy with a value over 0.50 and statistical significance level of Bartlett’s test below 0.05. A network analysis with the EBICglasso estimator, auto-correlation method and normalized centrality measures was performed to investigate the association among nonsocial cognitive skills and SC domains.

Finally, mediation analysis was performed in order to explore whether the alterations in SC were independently associated with the stage of dementia and AD severity (groups), or if others cognitive skills would mediate this response. Grouping variables in mediation analysis were represented as dummies of the clinical group compared to HC. A p-value<0.05 was used as a threshold to the level of significance. Jamovi and JASP software were used to perform statistical analysis.

## RESULTS

Sociodemographic and clinical characteristics of participants and differences among study groups are presented in Table 2. Age was higher in individuals with eAD and mAD compared to HC, while schooling was lower in eAD and mAD compared to HC. MMSE and CDR were progressively impaired in the MCI, eAD and mAD groups compared to HC group.

**TABLE 1.** Sociodemographic and clinical characteristics of participants by group.

|  | HC<br>*N=17* | MCI<br>*N=17* | eAD<br>*N=15* | mAD<br>*N=16* | p values |
| --- | --- | --- | --- | --- | --- |
| Age, years | 68.9 ± 6.67 | 71.4 ± 7.67 | 82.2 ± 4.87 *† | 77.4 ± 6.22* | < 0.001 |
| Schooling, years | 15 (13 - 16) | 12 (8 - 16) | 4 (4 - 11.5)* | 6 (4 - 14.3)* | 0.005 |
| MMSE | 28 (28-29) | 26 (26 - 28)* | 21 (20 - 23.5)*† | 15 (12 - 16)*†‡ | < 0.001 |
| CDR | 0 (0 - 0) | 0 (0 - 0.5)* | 1 (1 - 1.75)*† | 2 (1.5 - 2)*†‡ | < 0.001 |
| GDS | 3 (2 - 4) | 6 (4 - 8)* | 8 (5.5 - 10)* | 5.5 (2.75 - 8.25) | 0.003 |
| GAI | 3 (2 - 6) | 4 (2 - 8) | 5 (4 - 6) | 4 (2-8.25) | 0.777 |

Abbreviations: HC, Healthy controls; MCI, Mild Cognitive Impairment; eAD, early stage of Alzheimer's Disease; mAD, mild stage of Alzheimer's Disease; MMSE, Mini Mental State Examination; CDR, Clinical Dementia Rating; GDS, Geriatric Depression Scale; GAI, Geriatric Anxiety Inventory.
Age is presented as mean ± standard deviation, analyzed with one-way Analysis of Variance (ANOVA) test with post hoc de Tukey. The other variables are presented as median (25-75% quartiles), analyzed with Kruskal-Wallis's test.
\*p<0,05 in comparison to HC;

**TABLE 2 -.**
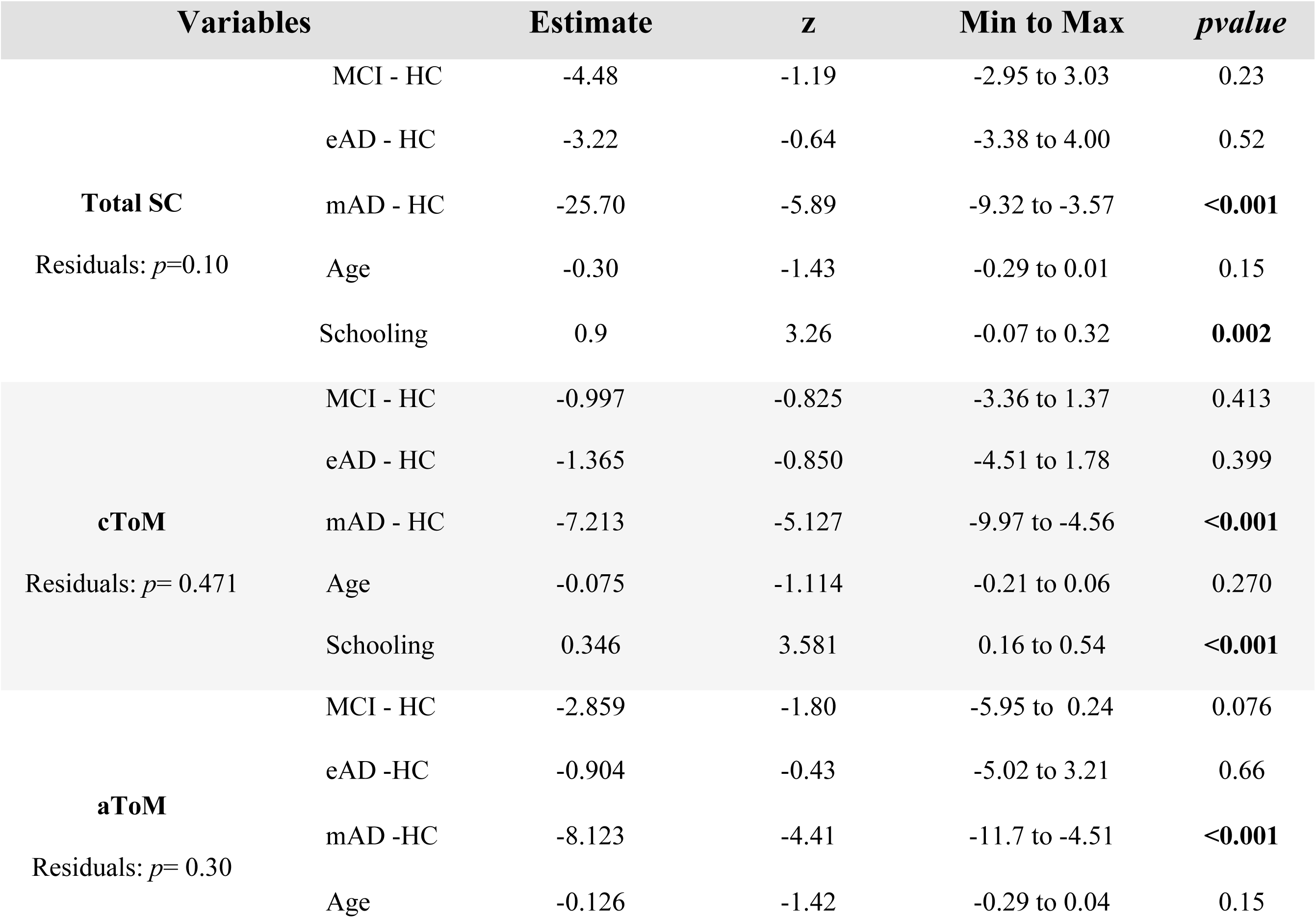
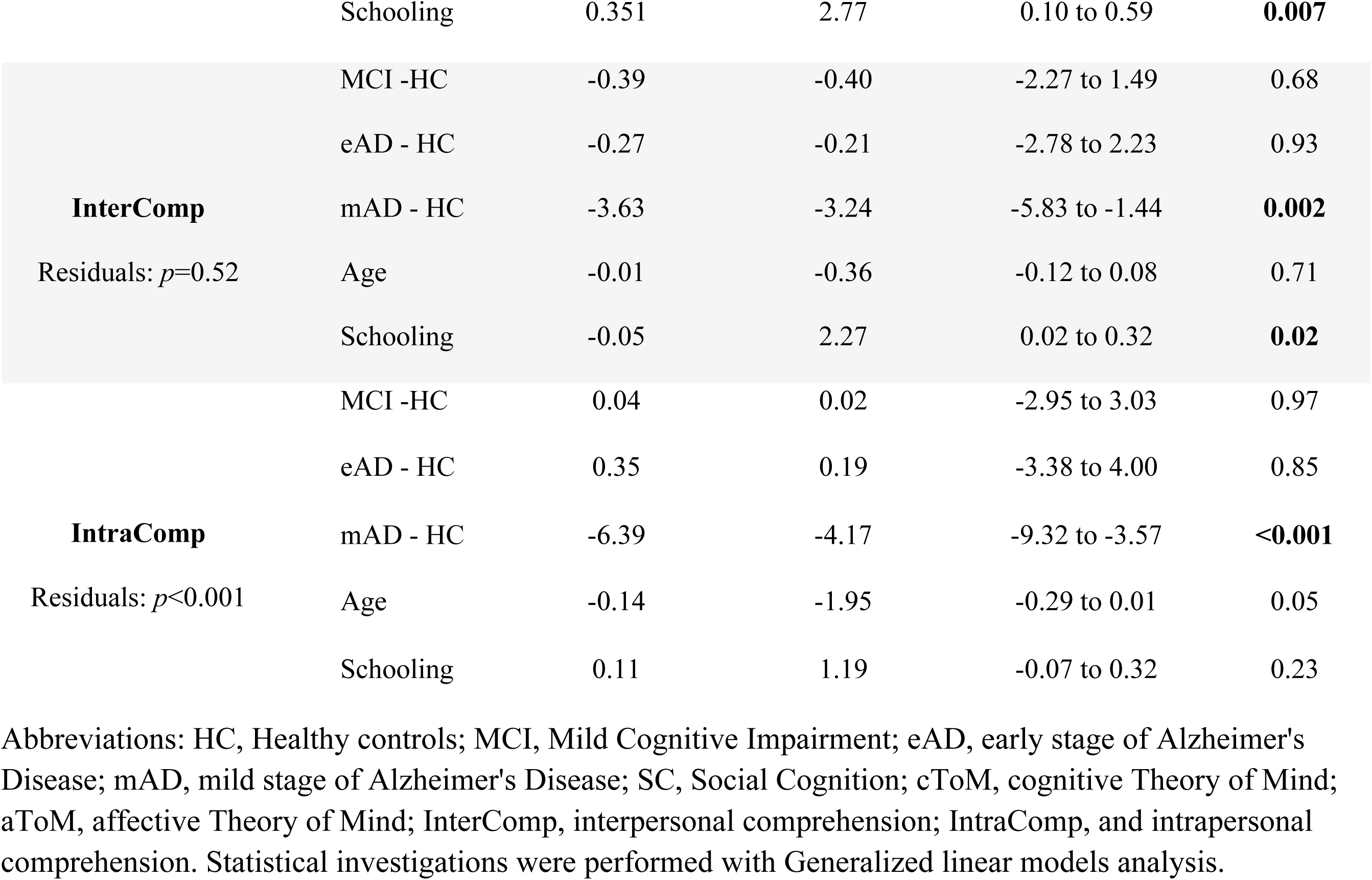
Effect of groups among Social Cognition domains, covariated by age and schooling.

### SC according to the severity of cognitive impairments

Since age and schooling were different among groups, these variables were inserted as covariates to prevent potential bias in the GzLM analysis investigating the effect of groups in SC. Years of schooling significantly influenced total SC, aToM, cToM and interpersonal comprehension, but not intrapersonal comprehension (see Table 2). Age did not show independent influence in any of the SC domains. All domains of SC were significantly affected by the group factor (Table 2, p<0.001).

The mAD (55.5 ± 16.3) group showed significantly lower scores of total SC compared to HC (88.9 ± 6.31), MCI (81.2 ± 9.81) and eAD groups (75.7 ± 11.8). Among the specific SC domains, the cToM was lower in the mAD (11.7 ± 5.39) compared to HC (21.4.9 ± 2.03) MCI (19.3 ± 3.10) and eAD (16.9 ± 3.81), as well as a lower aToM in the mAD (13.5 ± 5.42) compared with the other groups (HC 24.5+- 3.30, MCI 20.5 ± 5.39, eAD 19.8 ± 4.66). Moreover, the mAD group expressed both significantly impaired interpersonal comprehension (12.7 ± 3.30 mAD versus HC [17.4 ± 2.76], p<0.02, respectively), and intrapersonal comprehension compared to all three groups (17.6 ± 5.12 mAD versus HC 25.6 ±1.97, MCI 24.9 ± 2.00 and eAD 23.2 ± 3.71, p<0.02, respectively). Additionally, comparative analyses were executed to investigate possible effects of social appropriateness (socially adequate or inadequate) of videos on the participants’ performance according to the degree of severity, in each domain of SC. Results of social appropriateness were similar to overall scoring for each domain, except for the interpersonal comprehension in the videos of inadequate interactions. Groups were similar in respect of scoring for inadequate interaction in interpersonal comprehension domain (Mean Difference: HC versus MCI [-0.682; p=0.279], eAD [-0.808; p=0.335] and mAD [0.619; p=0.398]), therefore, this issue seems to be preserved even in the mAD group.

### Analyzing the links between nonsocial and social cognitive skills

Main nonsocial cognitive skills were resumed into factors as episodic memory, working memory, verbal fluency, verbal comprehension and executive functions (X^2^ = 13.7, p = 0.322, CFI = 0.996, TLI = 0.990, SRMR = 0.028; RMSEA = 0.046). Considering results from the principal component analysis as well as theoretical issues, episodic memory was assessed considering scores from both immediate and delayed recall of the logical memory test; working memory by both forwards and backwards digit span; verbal fluency from both semantic and phonological trials. Verbal comprehension (language) was assessed only by means of results in the WAIS-III comprehension subtest and executive functions by results on the clock drawing test.

Network analysis revealed 11 nodes, 35 non-zero edges and sparsity of 0.364 (see Figure 1). Verbal fluency, verbal comprehension and MMSE have the higher values of expected influence and strength of the network, followed by cToM and executive functions, denoting the importance of these variables. Furthermore, schooling and working memory have a negative influence on the network. Schooling seems to affect verbal comprehension, verbal fluency, executive functions and cToM.

**Fig. 1:**
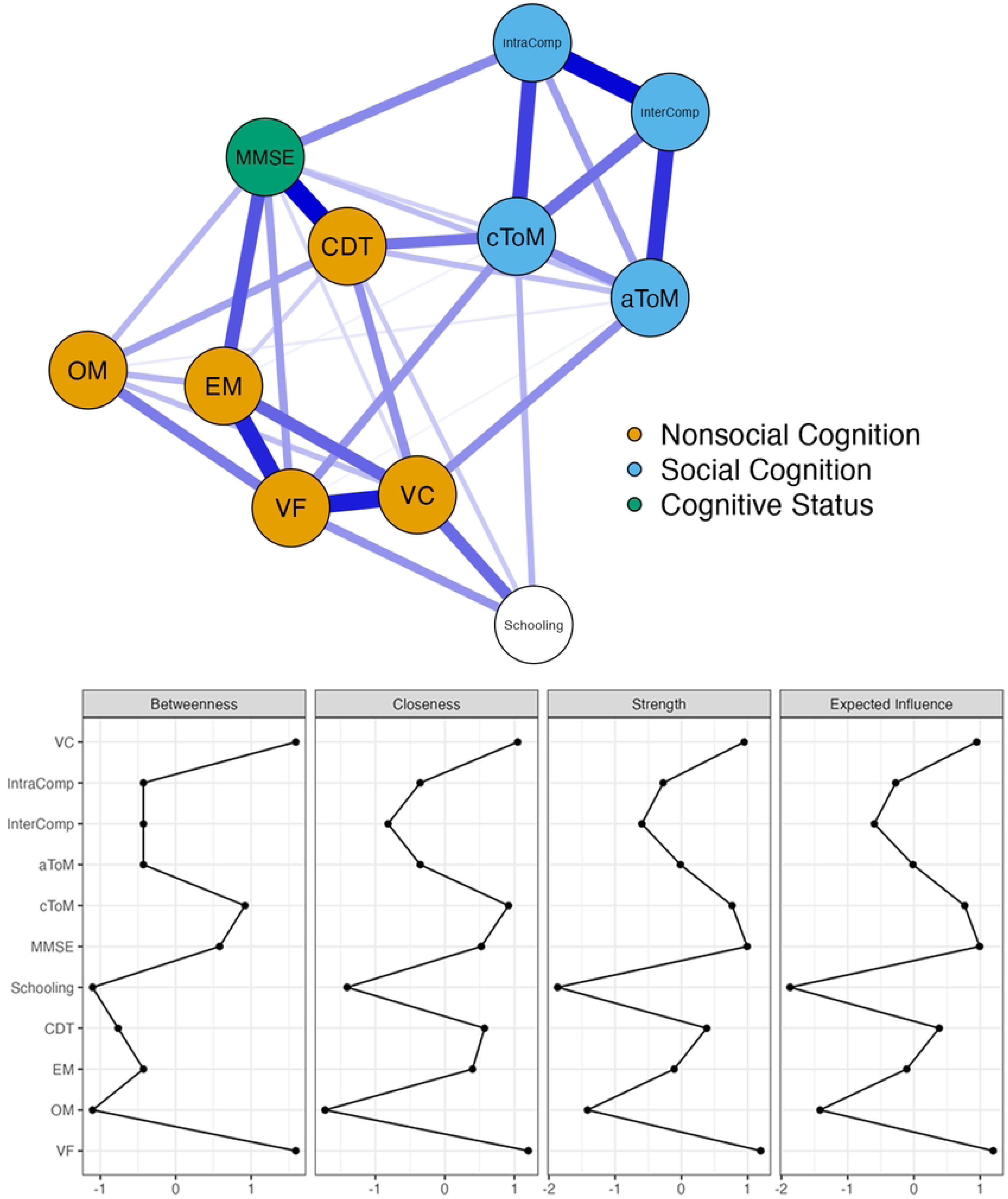
Network analysis with EBICGlasso between nonsocial and social cognitive skills, Mini-Mental State Examination (MMSE) and schooling.

SC domains were mainly associated with each other, which confirms and strengthens the specificity of these domains as a part of a particular cognitive skill. Comprehensions of social norms were not directly associated with any nonsocial cognitive skill. Whereas, ToM domains seems to be the bridge of SC with other nonsocial cognitive skills; since cToM presented connected paths with executive functions and verbal fluency and aToM connected with executive functions and verbal comprehension. More interestingly, SC domains exhibited no correlations with memory skills, including both working and episodic memory factors, thereby confirming the ESCOT as an assessment independent of memory influence (Figure 1).

By selectively investigating SC skills, a stronger association between MMSE and intrapersonal comprehension of social norms (IntraComp) was observed. MMSE was also associated with cToM and aToM. Additionally, MMSE, which was interpreted as a marker of dementia severity, revealed connections with all nonsocial cognitive skills, being the strongest connection with executive functions. By analyzing some of the triangular relations between variables, the findings of the network analysis may suggest a mediation effect of executive functions over cToM, such as a mediation effect of verbal fluency in cToM.

### Cognitive skills as mediators for SC

The mediation analysis revealed that episodic and working memory have no significant influence in the total SC (Figure 2A and 2B). Verbal fluency, verbal comprehension and executive functions were mediators of total SC score for all groups (Figure 2C, 2D and 2E). A direct effect of groups in the total SC score was observed only in the mAD (p<0.05). Verbal fluency, verbal comprehension and executive functions mediated the relation of groups with the total SC (Figures 2C, 2D and 2E).

**FIG. 2:**
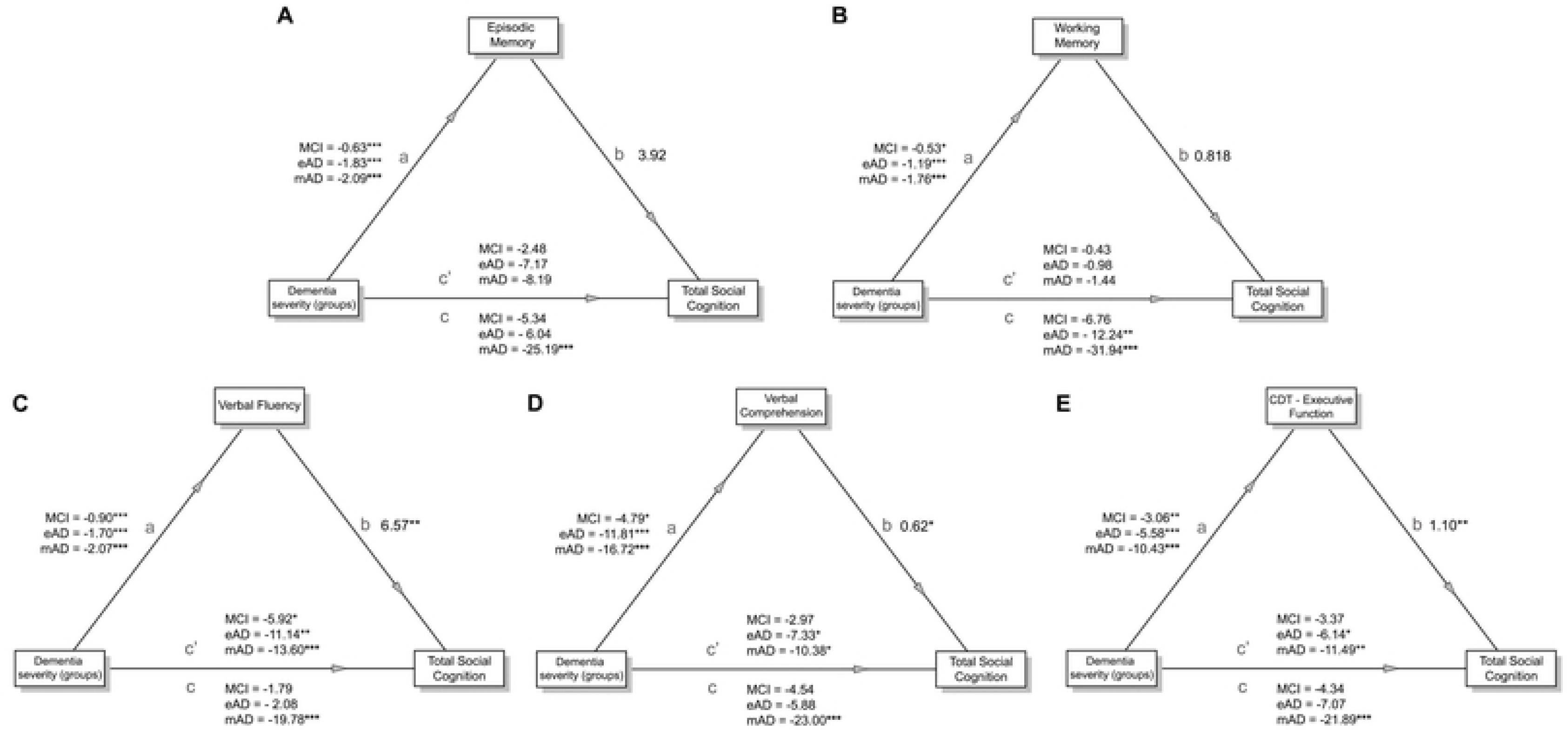
Path diagrams of mediation analyses investigating cognitive factors as possible mediators of dementia severity groups in total SC. The coefficients are presented for each pathway, where (a) reveals the association of groups with the cognitive factor, (b) reports the influence of cognitive factor with total SC, (c) expresses the direct effect and (c’) is the indirect pathway mediated by the cognitive skill factor. A. Episodic memory; B: Working memory; C: Verbal fluency; D: Verbal comprehension; E: Executive functions.

Since verbal fluency, verbal comprehension and executive functions mediated the relation of mAD with total SC, an investigation of the specific domains of SC was performed separately. In the GzLM analysis, schooling was significantly associated with ToM and interpersonal comprehension domains, thus, the mediation analysis was later adjusted for this variable. Given the small sample size, a bootstrap estimation was used with 1000 replications to ensure that results found in the mediation analysis covariated by schooling.

Verbal fluency seemed to exert an indirect effect in the cToM among all groups (supplementary material). However, when the mediation analysis was covariated by the schooling, the effects persisted significantly only in the mAD group comparison (Estimate = −2.70, CI95% −5.35 to −0.04, p=0.046, Figure 3), but not in the eAD (Estimate = −2.05, CI95% −4.10 to 0.01, p=0.051) neither MCI (Estimate = −1.15, CI95% −2.39 to 0.08, p=0.068). Moreover, aToM seemed indirectly affected by verbal fluency among all groups, but when the mediation analysis was covariated by the schooling, the mediation effects did not persist significantly (Figure 3, Suppl). No indirect effect of verbal fluency was observed in the intrapersonal and interpersonal comprehension domains (p>0.05, Figure 3A). Only the mAD group presented a direct effect in the investigations of verbal fluency (p<0.03).

**FIG. 3:**
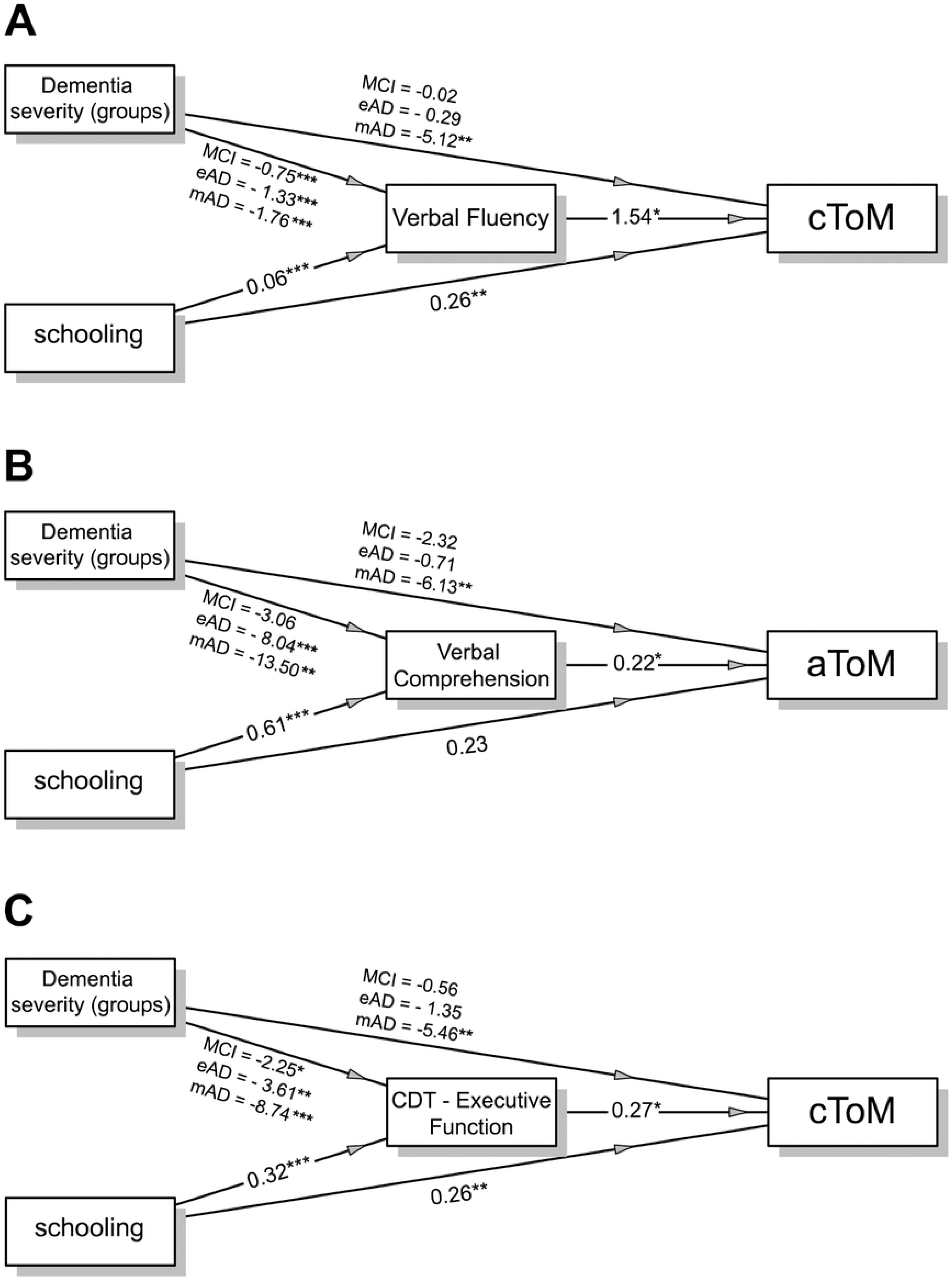
Path diagrams of mediation analyses investigating cognitive factors as possible mediators of dementia severity groups in specific domains of SC with schooling as a covariate variable. The coefficients are presented for each pathway with each dummy for clinical groups comparison with Healthy Controls. A. Verbal fluency; B: Verbal comprehension; C: executive functions.

Verbal comprehension had a mediation effect in the cToM and aToM among groups (Supplementary material). Nevertheless, by introducing the schooling as covariate, the mediation effects did not persist statistically significant in cToM (MCI: Estimate = −0.13, CI95% −1.07 to 0.31, p=0.69; eAD: Estimate = −0.34, CI95% −1.89 to 1.03, p=0.63; mAD: Estimate = −0.57, CI95% −2.81 to 1.67, p=0.63). With respect to aToM, the mediation effects persisted significantly in the mAD (mAD: Estimate = −2.97, CI95% −6.43 to −0.38, p=0.044, Figure 3B), but not for MCI and eAD groups (MCI: Estimate = −0.67, CI95% −2.17 to 0.02, p=0.20; eAD: Estimate = −1.77, CI95% −4.75 to −0.29, p=0.08). Therefore, verbal comprehension exerted an indirect effect on aToM only in mAD patients. Intrapersonal and interpersonal comprehension was not influenced by verbal comprehension. A direct effect in SC domains was observed in the mAD in the verbal comprehension analysis.

Lastly, executive functions exerted a mediation effect in cToM and aToM among groups in the simple mediation analysis. When schooling was introduced as covariate, the executive functions only persisted as a mediator of cToM in the mAD (Figure 3C, mAD: Estimate = −2.37, CI95% −4.61 to −0.12, p=0.038 versus eAD: Estimate = −0.98, CI95% −2.08 to 0.13, p=0.08; MCI: Estimate = −0.61, CI95% −1.41 to −0.20, p=0.14 . Comprehensions of social norms were neither not related to, nor influenced by executive functions (p>0.05, Supplementary Material).

## Discussion

The main results of the current study revealed global alterations in SC only in the moderate stage of AD after controlling for confounder factors. The neuropsychological battery chosen for assessment of SC were proved to be independent of the influence of working and episodic memory. Nevertheless, specific domains of SC (more precisely ToM) were indirectly influenced by verbal fluency, verbal comprehension and executive functions.

An initial sample characterization revealed that patients in the AD groups presented older age and lower schooling compared to the HC group. This result is somewhat expected and reflects an inherent characteristic of pathological aging, since age is the greatest risk factor known for AD manifestation ^22^. Moreover, lower schooling implies lower cognitive reserve, recognized as a protective factor for dementia ^23^. Since age and schooling could act as confounding factors for SC performance, our analysis controlled for these variables to prevent bias. Results revealed an effect of schooling, but not age, over specific domains and overall SC. This finding suggests that social environment variables influencing individuals’ experiences that enhance understanding and vocabulary on emotions, empathy and social rules may be more relevant than biological characteristics of aging in preserving SC skills.

In fact, global SC were impaired only in the moderate stage of AD when age and schooling were taken into account. This finding may help to tackle controversies with respect to the stage of the disease in which SC impairments emerge. Some studies have found that such impairments are already present in the early stage ^11^, while in others it was revealed only in the moderate and severe stages ^12^. Lack of control of sample characteristics in baseline, such as schooling degree, may be one of the main reasons for these differences regarding time to onset SC deficit. Another possible explanation for controversial findings in literature is related to the complexity of neuropsychological instruments used for assessment of SC. For example, if the task demands retention of stimuli, memory skills may prevent a more independent assessment of SC tasks. Fliss and collaborators kept visible stimuli during their battery of tasks to reduce working memory effects, in a way similar to ours^12^ . The network analysis of the present study revealed that SC skills did not correlate with working and episodic memory factors, certifying the ESCOT as an instrument that mitigates the influence of memory.

With regard to investigation of selectivity of SC dysfunctions, our results showed that both cognitive and affective ToM domains were impaired only among mAD participants. These findings are in consonance with those reported in Demichelis meta-analysis ^24^, according to which both cognitive and affective ToM are impaired in AD compared with HC. Therefore, ToM may be preserved until an initial phase of dementia. Complaints about ToM problems could be therefore an early cognitive marker of the evolution of the disease. Impairments in interpersonal and intrapersonal understanding of social norms were also observed only in the moderate stage group. In the study by Asaad et al (2018) with healthy elderly participants, only interpersonal understanding of social rules changed with age.

Difficulties in understanding social norms may be due to language impairments. For example, Halberstadt and colleagues ^25^ in a sample of 60 young adults aged 36 to 59 years old and 61 older adults aged 60 to 85 years old, observed a smaller vocabulary of older adults to discriminate social behaviors as appropriate and inappropriate when they saw short videos with social interaction. Our results, therefore, contradict these findings from the literature by identifying that interpersonal and intrapersonal comprehension skills of social norms are preserved until an early stage of AD. Despite this, one should consider the influence of schooling or the increase of narrative weaknesses with age. Our results suggest that schooling is a significant predictor for SC losses.

Another aspect to be highlighted is that participants from the mild AD group retained their interpersonal comprehension of social norms skills only in the inappropriate interaction videos. These findings may be related to the fact that emotional content is more evident in videos of inappropriate interaction as they highlight everyday situations that violate socially established norms. Perrim and colleagues ^26^ identified that emotional content benefited long-term memory in patients in the early phase of AD. This occurs because emotional memory activates the amygdala-hippocampal region that occurs in a broader frontotemporal network. The amygdala is primarily involved in the essential memory of emotional messages. Studies also show that new negative emotional content can remain preserved as the disease progresses ^27^ .

The network analysis revealed SC domains closely associated with each other, which reinforces that SC is a cognitive skill isolated from nonsocial cognition. Among SC skills investigated in this study, our findings suggest that ToM is the bridge for the interactions with nonsocial cognitive functions. Possible explanations may be considered from a neurodevelopmental perspective. ToM is an ability acquired in early life. On the other hand, comprehension of social norms results predominantly from social learning, as a consequence of the continuous exposition to experiences in an social and cultural organized environment. Therefore, selective losses in social norms comprehension should be considered as a clinically relevant marker for aggravation of AD. This possibility is reinforced by the preservation of understanding social norms as observed in older healthy elderly in the original study^8^. To the best of our knowledge, this study is the first to explore the comprehension of social norms according to the AD stage, highlighting the uniqueness of this ability to detect SC impairment in pathological aging.

Additionally, comprehension of social norms was not mediated by any nonsocial cognition skills. On the other hand, ToM was mediated by verbal fluency, verbal comprehension and executive functioning, as investigated by the CDT. By adjusting the mediation effect by the possible confounder (i.e. schooling), we identified that selective domains of ToM were influenced by specific factors. Cognitive ToM was mediated by verbal fluency and executive functioning, while affective ToM was mediated only by verbal comprehension. Both results were independent of level of schooling. Current studies have reported that ToM is a multidimensional function that involves distinct brain networks and therefore explain neuropsychological dissociations ^28^ . The insular cortex is more associated with aToM while the prefrontal cortex is associated with cToM. Our findings that only cToM was mediated by Executive Functions may be explained by differences in the activation of these brain networks. Furthermore, judging what the other is thinking depends on greater controlled processing, whereas, in addition to understanding the story, it requires greater abstraction of the other’s internal state in the interaction context. Therefore, our results indicate that abstraction impairments, in the moderate stage of AD, mediate the ability to interpret other’s thoughts, in other words, cToM.

On the other hand, judging what the other is feeling (aToM) involves more automatic processes due to emotional contagion. Our findings that aToM was mediated only by verbal comprehension may be associated with the difficulty in understanding the other’s emotional state in a static situation, since in this case, controlled processing is required. Although ESCOT is an ecological test, it is understood that, for individuals in a moderate phase of AD, procedures in the context of real interaction would be more appropriate to access automatic emotional ToM processes. In a study involving ToM assessments in a real social context, changes in cToM and aToM were identified in a sample of 20 participants in the early stages of AD ^29^.

Regarding the influence of verbal fluency, It must be considered that the evaluation of performance in the ESCOT requires complex verbal responses that demand interpretation and judgment of the social situation, in addition to lexical access and organization of the narrative. To be classified with a maximum score (3 points), a cToM response had to associate a social content with a contextual reason. For example, in the situation where an elderly woman’s bag breaks and a young man passes by without helping, the maximum scoring response would be something like “She thought he would help, because her bag ripped and she certainly has difficulty collecting her belongings”. An answer with only social content, such as “she thought he would help her” would result in a sum of 2 points. In other words, apparently in AD, impairments in verbal fluency and restricted vocabulary influence the ability to present a comprehensive answer about the other’s mental states. Future studies, therefore, can better assess Theory of Mind in AD if procedures are connected to real contexts with less requirement for verbal fluency and complex responses.

In sum, the ESCOT test enabled the assessment of Social Cognition mitigating the influence of memory-skills in the evolution of Alzheimer’s Disease severity. By adjusting for age and schooling, deficits in ToM and comprehension of social norms were observed only in the moderate stage of Alzheimer’s Disease. Social norms comprehension for negative situations seems preserved even in the moderate stage of Alzheimer’s Disease. Performance in cognitive ToM was influenced by executive function and verbal fluency and Affective ToM was moderate by verbal comprehension. It is suggested that CS deficits can be indicators of AD severity and the identification of changes in CS skills can contribute to the differential diagnosis and help in directing neuropsychological rehabilitation. Knowing that socio-cognitive training is effective in improving ToM skills ^30^, identifying declines in CS in healthy elderly people or those in the early stages of dementia can enable interventions to stimulate these skills and provide greater quality of social interaction for this population. Furthermore, considering that the understanding of social norms proved to be an independent measure of non-social cognitive abilities, the assessment of this SC component proved to be an important measure to evaluate CS in individuals with cognitive impairments. Changes in this ability may prove to be an early marker for the disease.

## Acknowledgements.

This study was financed in part by the Coordenação de Aperfeiçoamento de Pessoal de Nível Superior – Brasil (CAPES) – Finance Code 001. The authors are grateful for the continued institutional and financial support from Associação Fundo de Incentivo à Pesquisa (AFIP) that has made possible the realization of clinical assistance and research.

## Author contributions

Conceptualization: G.C.M.,V.L.D.V., C.B.M. Investigation: G.C.M, V.L.D.V, F.M.P, A.M.L.A, D.M.S, Methodology: G.C.M.,V.L.D.V., C.B.M., F.M.P, A.M.L.A, D.M.S, I.S.R., Validation: C.B.M., I.S.R., P.H.O.B., S.T. Writing-original draft preparation.G.C.M.,V.L.D.V., C.B.M., F.M.P, A.M.L.A, D.M.S,I.S.R. Writing-review and editing. G.C.M; C.B.M., I.S.R., P.H.O.B. S.T.

